# Control of microtubule dynamics using an optogenetic microtubule plus end–F-actin cross-linker

**DOI:** 10.1101/142414

**Authors:** Rebecca C. Adikes, Ryan A. Hallett, Brian F. Saway, Brian Kuhlman, Kevin C. Slep

## Abstract

We developed a novel optogenetic tool, SxIP-iLID, to facilitate the reversible recruitment of factors to microtubule (MT) plus ends in an End Binding (EB) protein-dependent manner using blue light. We show that SxIP-iLID can track MT plus ends and recruit tgRFP-SspB upon blue light activation. We then used this system to investigate the effects of cross-linking MT plus ends and F-actin in *Drosophila* S2 cells to gain insight into spectraplakin function and mechanism. We show that SxIP-iLID can be used to temporally recruit a F-actin binding domain to MT plus ends and cross-link the MT and F-actin networks. Light-mediated MT-F-actin cross-linking decreases MT growth velocities and generates a MT exclusion zone in the lamella. SxIP-iLID facilitates the general recruitment of specific factors to MT plus ends with temporal control enabling researchers to systematically regulate MT plus end dynamics and probe MT plus end function in many biological processes.

**Summary:** SxIP-iLID is a novel optogenetic tool designed to assess the spatiotemporal role of proteins on microtubule dynamics. We establish that optogenetic cross-linking of microtubule and actin networks decreases MT growth velocities and increases the cell area void of microtubules.

## Introduction

Cellular and developmental processes require the spatial and temporal control of protein-protein interactions. The cytoskeleton, composed of MTs, F-actin, and intermediate filaments, is tightly regulated and remodeled throughout the cell cycle. How proteins regulate cytoskeletal dynamics and mediate cross-talk between the networks is an active area of research. For example, the dynamic coupling of the actin and MT networks is essential for neuronal growth (Tortosa et al., 2011, Sanchez-Soriano et al., 2009, Lee et al., 2000; Lee and Luo, 1999; Prokop et al., 1998), cell shape changes, migration (Guo et al., 1995, Wu et al., 2008, Wu et al., 2011) and determining the site of the contractile ring (Kunda and Baum, 2009),.

Historically, probing the role of protein-protein interactions in complex cellular networks with spatial and temporal resolution has been difficult. However, recent advances in cellular optogenetic techniques have enabled biologists to dissect the spatiotemporal mechanisms that regulate diverse cellular systems. Many inducible protein dimer systems have recently been generated, optimized, and made available to researchers to control protein activity and/or localization within cells and organisms. Currently available dimer systems include both chemically induced dimers, such as the FRB/FKBP12 system that can be heterodimerized with rapamycin (Rivera et al., 1996), and light inducible dimers (LIDs). LIDs come from photoactivatable systems naturally occurring in plants and allow for regional, reversible activation using light. LIDs include phytochromes, cryptochromes, and Light-Oxygen-Voltage (LOV) domains. LOV domains have been used in engineered homodimer paired systems such as tunable light-controlled interacting protein tags (TULIPs (LOVpep/ePDZb)) (Strickland et al., 2012), improved light-inducible dimer (iLID) (iLID/SspB) (Guntas et al., 2015), and Zdk/LOV2 – a heterodimer that dissociates when photoactivated (Wang and Hahn, 2016). LOV-based systems rely on a blue light-dependent conformational change in the LOV2 domain that facilitates the release and unfolding of an α-helix termed the Jα helix. The iLID/SspB system contains a short ssrA peptide sequence embedded in the Jα helix of the LOV domain. The ssrA sequence is occluded from binding its partner SspB in the dark. However, upon blue light activation, the ssrA sequence becomes accessible and can bind SspB. Specific advantages of the iLID/SspB system include 1) no off-target effects in nonplant eukaryotes, and 2) the availability of a suite of iLID constructs with different on/off kinetics and SspB binding affinities (Guntas et al., 2015, Hallett et al., 2016, Zimmerman et al., 2016).

iLID as well as other LIDs have been used to perturb pathways involved in cell protrusion (Hallett et al., 2016), and cell migration (Weitzman and Hahn, 2014), to activate formins to control actin architecture (Rao et al., 2013), and regulate organelle transport and positioning (van Berjeijk et al., 2015, Duan et al., 2015). Most recently the Zdk/LOV2 system was used to dissociate the MT plus end protein EB1 with temporal and spatial control. This study revealed that the equilibrium of MT polymerization dynamics changes in under a minute and the MT network rapidly reshapes (van Haren et al., 2017 *Preprint*). However, a system to recruit select proteins or protein domains of interest to MT plus ends has not been created.

Here, we develop and validate an optogenetically controlled MT plus end recruitment system, which can be easily adapted to answer a variety of questions pertaining to the regulation of MT dynamics and MT network organization. To create this system we utilized iLID (iLID/SspB). This system is ideal for the generation of reversible recruitment of specific SspB-fusion proteins to MT plus ends. Here, we show that EB-binding SxIP motifs (Slep et al., 2005; Honnappa et al., 2009), appended to the iLID N-terminus, confer iLID with MT plus end tracking activity. When exposed to blue light, SxIP-iLID recruits SspB-tagged proteins to MT plus ends. We establish SxIP-iLID as a tool with broad utility that can be used to systematically study the mechanisms of MT regulators and MT associated proteins.

We then utilized SxIP-iLID to investigate how MT-F-actin cytoskeletal cross-linking affects MT dynamics and cell morphology. Previous studies have shown that coordination between F-actin and MTs is important for cell migration, mitosis, and tissue morphogenesis. One class of cytoskeletal cross-linkers are the spectraplakins. Spectraplakin loss of function leads to severe axon shortening and MT disorganization (Sanchez-Soriano et al., 2009). Mutations in the *Drosophila* actin-MT cross-linking protein Shot cause a variety of cellular and tissue defects including changes in actin-MT organization, cell-cell adhesion, and integrin mediated epidermal attachments to muscle (Gregory and Brown, 1998; Walsh and Brown, 1998; Propkop et al., 1998; Strumpf and Volk 1998; Röper and Brown, 2003). Conditional knock out of the spectraplakin Actin Cross-linking Factor 7 (ACF7) in mice yields defects in cell migration (Wu et al., 2008, Goryunov et al., 2010). These mutational and knock out experiments provide information on long-term whole tissue depletion of a spectraplakin, however having a subcellular temporal and rapidly reversible way to probe the effects of cross-linking will provide mechanistic details on the direct cellular changes induced by cross-linking. Proteins of the spectraplakin family typically contain two N-terminal calponin-homology (CH)-type F-actin binding domains, and a C-terminal MT-binding module consisting of a Gas2-related (GAR) domain, Gly-Ser-Arg rich (GSR) region, and an EB-binding Sx(I/L)P motif (Lee et al., 2000; Slep et al., 2005, Wu et al., 2008; Applewhite et al., 2010). Although recent studies have proposed mechanisms for the regulation of spectraplakins (Wu et al., 2011; Kapur et al., 2012; Applewhite et al., 2013; Takács et al., 2017), the direct downstream cellular outputs of regulated cross-linking remain poorly understood. To begin to understand how cross-linking affects cytoskeletal dynamics and network organization we utilized the SxIP-iLID system to optogenetically cross-link MTs and F-actin. We show that light-mediated MT-actin cross-linking decreases MT growth velocities and creates a MT exclusion zone in the lamella.

## Results

### Design of a light-inducible system for MT plus end tracking

Our goal was to control the spatial and temporal recruitment of proteins to the MT plus end. To do so we designed a switch that would constitutively track MT plus ends and recruit a protein or domain of interest upon blue light activation (Fig. 1, A and B). To generate a MT plus end recruitment switch we capitalized on the ability of the EB MT plus end tracking protein family to directly bind cellular factors that contain a SxIP motif (Slep et al., 2005; Honnappa et al., 2009). We fused individual or arrayed EB-binding SxIP motifs to the iLID domain. We designed four SxIP-iLID constructs to generate a library with a range of MT plus end tracking activity. All constructs contained a N-terminal eGFP to allow detection and tracking (Fig. 1A). The first construct, SKIP-iLID, contains a single 18-amino acid SKIP motif from the spectraplakin MT-actin crosslink factor 2 (MACF2). Two constructs contain a pair of tandemly arrayed SxIP motifs, either SKIP-SKIP or SKIP-SRIP, with the two motifs separated by the linker sequence that bridges the two endogenous SxIP motifs in cytoplasmic linker protein-associated protein 2 (CLASP2) (Honnappa et al., 2009). A fourth construct contains a single MACF2 SKIP motif followed by the GCN4 leucine zipper (LZ) homodimerization domain (Fig. 1A and Table 1)(Honnappa et al., 2009, Steinmetz et al., 2007). The tandem SKIP constructs and the dimerized SKIP construct were designed to enhance binding to EB dimers via avidity, which has previously been shown to enhance the apparent MT plus end tracking activity of SxIP constructs (Honnapppa et al., 2009).

**Figure 1.**
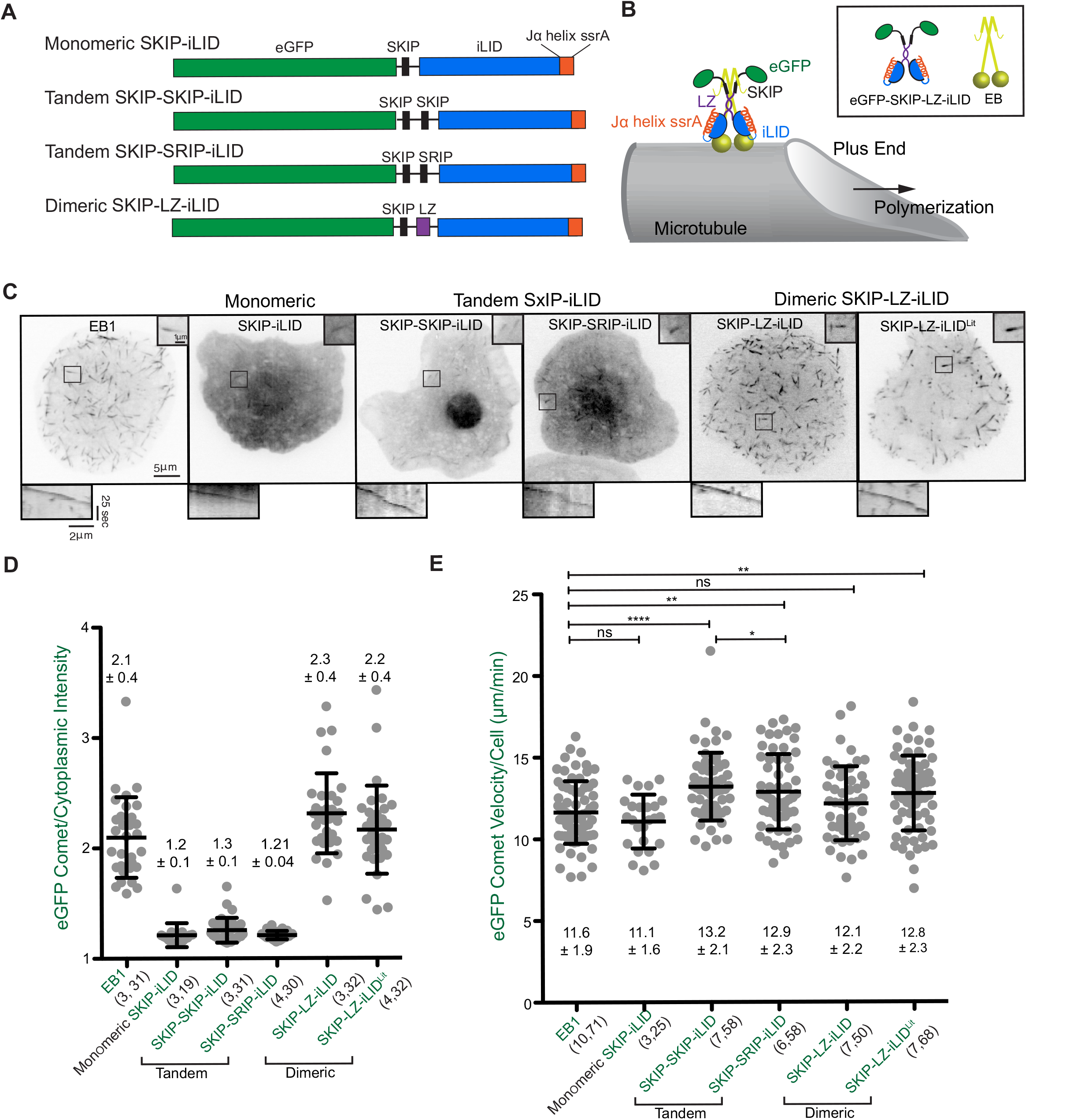
Figure 1. **SxIP-iLID constructs track MT plus ends and do not dramatically perturb MT comet velocities. (A)** Schematic of eGFP-labeled SxIP-iLID constructs. **(B)** Cartoon diagraming eGFP-SKIP-LZ-iLID at a MT plus end. **(C)** Representative images of eGFP-SxIP-iLID constructs in *Drosophila* S2 cells. Scale bar, 5 μm, inset scale bar, 1μm. Comets can be observed in all cells expressing an eGFP-SxIP-iLID construct. Kymographs below each image show a representative eGFP-SxIP-iLID MT plus end comet. Scale bars, 2 μm, 25 seconds (see accompanying Video 1). **(D)** The ratio of mean eGFP-SxIP-iLID construct intensity per pixel on a MT plus end relative to the mean intensity per pixel in an adjacent cytoplasmic region. The dimeric SKIP-LZ-iLID construct shows the greatest ratio of eGFP comet intensity relative to cytoplasmic background, comparable to that calculated for EB1-GFP. Error bars indicate the SD. Numbers in parenthesis indicate (number of experiments, total number of cells quantified). **(E)** SxIP-iLID constructs show some variation in eGFP comet velocities, however, the mean comet velocity of the dimeric SKIP-LZ-iLID construct is not significantly different from the mean comet velocity of an EB1-GFP control. Error bars indicate the SD. Numbers in parenthesis indicate (number of experiments, total number of cells quantified). P-values were determined by two-way unpaired Student’s t-test. * p<0.05, ** p<0.005, **** p<0.0001.

**Table 1:**
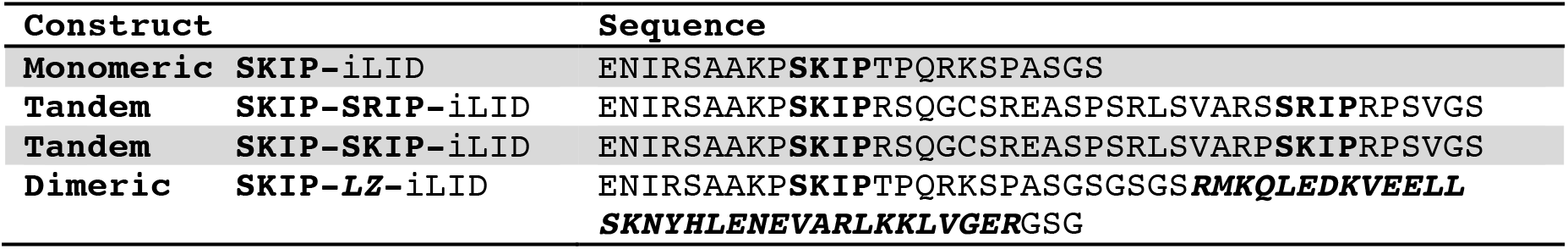
Sequences of SxIP Motifs, Linkers, and Leucine Zipper for iLID constructs

We assayed the ability of these constructs to track polymerizing MT plus ends in *Drosophila* S2 cells using time-lapse eGFP imaging. Each of the four SxIP-iLID constructs exhibited MT plus end tracking activity as revealed by distinctive comet-like patterns that moved throughout the cytoplasm (Fig. 1C and Video 1). Monomeric SKIP-iLID comets were dim and difficult to observe in most cells, while the dimeric SKIP-LZ-iLID exhibited robust MT plus ends tracking activity (Fig. 1C and Video 1). Ratiometric analysis of the eGFP comet:cytoplasmic intensities revealed that the levels of monomeric and tandem SxIP-iLID constructs on MT plus ends were only slightly enhanced above their respective cytoplasmic level (~1.2x) whereas the level of the dimeric SKIP-LZ-iLID construct on MT plus ends was 2.3 times higher than cytoplasmic, similar to the ratio measured for EB1-GFP (Fig. 1D). While the monomeric and tandem SxIP-iLID constructs exhibited varying degrees of nuclear localization, the SKIP-LZ-iLID construct showed minimal nuclear localization. As a test tool, we generated a constitutively active, lit mimetic construct, SKIP-LZ-iLID^Lit^ (iLID^I539E^)(Harper et al., 2004, Lungu et al., 2012), that is competent to bind SspB without blue light activation. SKIP-LZ-iLID^Llt^ robustly tracked MT plus ends (Fig. 1C and Video 1).

To determine if SxIP-iLID constructs altered MT plus end polymerization rates, we tracked individual eGFP-SxIP-iLID comets over time. There was no significant difference between the EB1 control and the monomeric SKIP-iLID construct, nor the dimeric SKIP-LZ-iLID WT or lit mimetic constructs, however, there was a slight increase in the average comet velocities for the tandem SxIP-iLID constructs (Fig. 1E).

To determine if SxIP-iLID construct expression level, or the amount of SxIP-iLID construct at MT plus ends affected comet velocities we analyzed the total cell intensity/cell area and the MT comet/cytoplasmic intensities in each cell and plotted this against the average comet velocity. To assess the strength of the correlation we determined the Pearson’s correlation coefficient (Fig. S1, A and B). Using this statistical assessment we established that the only construct which showed a slightly significant correlation between total cell intensity or plus end fluorescence intensity with comet velocity was the tandem SKIP-SRIP-iLID construct (total cell intensity/cell area p=0.017 and MT comet/cytoplasmic intensity p=0.036).

This establishes a library of SxIP-iLID constructs with differential MT plus end tracking activities and highlights the interesting observation that specific, multivalent EB-binding scaffolds can themselves affect (enhance) MT growth rates.

### The SxIP-iLID switch recruits tgRFP-SspB upon photactivation

To determine whether the SxIP-iLID constructs could recruit an SspB-tagged protein to polymerizing MT plus ends we observed localization of a tgRFP-SspB construct pre and post photoactivation (Fig. 2, A-C; and Video 2). We transfected *Drosophila* S2 cells with a SxIP-iLID construct (either a tandem SxIP-iLID construct or the SKIP-LZ-iLID construct) as well as tgRFP-SspB (Fig. 2A). Prior to blue light photoactivation, tgRFP-SspB had a diffuse cytoplasmic signal (Fig. 2C). Strikingly, upon blue light photoactivation, tgRFP-SspB rapidly colocalized with the SxIP-iLID constructs at polymerizing MT plus ends (Fig. 2C). This demonstrates that blue light can effectively liberate the SsrA sequence from its dark state embedded conformation in the iLID Jα helix, to an open conformation competent to bind and recruit an SspB-tagged protein to the MT plus end. As observed with the SxIP-iLID modules, more robust tgRFP-SspB MT plus end recruitment was observed with the dimeric SKIP-LZ-iLID construct (Fig. 2C and Video 2). In comparison, the constitutively active lit mimetic (SKIP-LZ-iLID^Lit^) recruited tgRFP-SspB to MT plus ends both pre- and post-blue light exposure, demonstrating the efficacy of ssrA-SspB engagement (Fig. 2C). To determine if recruitment of tgRFP-SspB to MT plus ends altered comet velocities we continuously activated SxIP-iLID constructs with pulses of blue light, thereby maintaining SxIP-iLID - tgRFP-SspB interaction and tgRFP-SspB MT plus end localization over time. Using the eGFP signal from the SxIP-iLID constructs, we again acquired time-lapse movies and used MTrackJ to determine comet velocities (Fig. 2D). tgRFP-SspB MT plus end recruitment via dimeric SKIP-LZ-iLID yielded comet velocities that were not statistically different from control cells containing tgRFP-SspB and EB1-GFP and no SxIP-iLID construct. Continually activated tandem SxIP-iLID constructs also recruited tgRFP-SspB to MT plus ends, and did not alter comet velocities as compared to the tandem SxIP-iLID construct alone (Fig. 2D). Monomeric SKIP-iLID construct plus end tracking activity was too weak to ascertain tgRFP-SspB MT plus end recruitment and was not pursued further. This establishes a two-component blue light-inducible heterodimerization system for recruiting proteins of interest to MT plus ends with temporal resolution.

**Figure 2.**
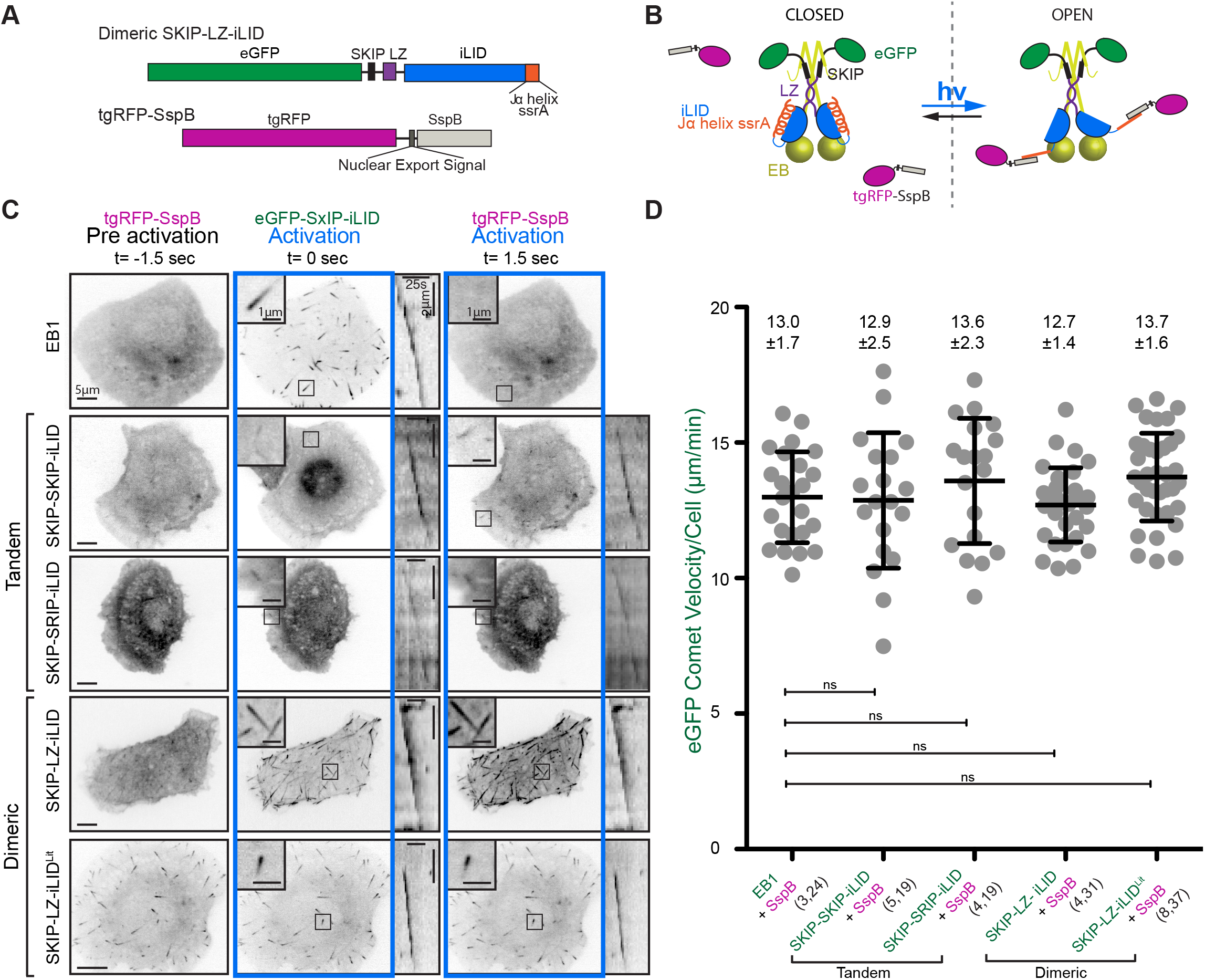
Figure 2. **Photoactivated SxIP-iLID constructs rapidly recruit tgRFP-SspB to MT plus ends without altering MT comet velocities. (A)** Diagram of SKIP-LZ-iLID and tgRFP-SspB constructs. **(B)** Cartoon diagraming the binding of tgRFP-SspB to SKIP-LZ-iLID upon blue light activation (hv). SKIP-LZ-iLID localizes to MT plus ends via association with EB dimers. When exposed to blue light the iLID Jα helix releases from the LOV domain. The ssrA motif embedded in the Jα helix becomes accessible to bind SspB and thereby recruits tgRFP-SspB to MT plus ends. **(C)** Representative images of S2 cells co-transfected with either a tandem SxIP-iLID construct or a dimeric SKIP-LZ-iLID construct and a tgRFP-SspB control construct, repeatedly pulsed with blue light (250 msec) every 3 seconds at 488 nm. Representative images are of tgRFP-SspB localization pre and post photoactivation in *Drosophila* S2 cells (see accompanying Video 2). The SKIP-LZ-iLID^Lit^ construct is constitutively competent to bind SspB in the absence of blue light. Scale bars at left, 5 μm, inset scale bar, 1 μm. Kymograph scale bars: 2 μm, 25 seconds. **(D)** SxIP-iLID-based recruitment of tgRFP-SspB to MT plus ends does not significantly alter mean MT plus end comet velocities compared to control cells co-transfected with EB1-GFP and tgRFP-SspB. Error bars indicate SD. Numbers in parenthesis indicate (number of experiments, total number of cells quantified). P-values were determined by two-way unpaired Student’s t-test.

### tgRFP-SspB MT plus end recruitment is rapidly reversible

In order for an optogenetic system to regulate cytoskeletal dynamics with spatial and temporal efficacy, the system must transition to an active or inactive state within seconds. We observed rapid (< 1.5 second) recruitment of tgRFP-SspB to MT plus ends that peaks between 12 and 33 seconds post activation (Figs. 2C, 3A and B; and Video 3). The decay of tgRFP-SspB signal from MT plus ends post activation reflects an apparent dissociation rate. This rate is a convolution of many factors including the nucleotide state of tubulin at growing MT plus ends, the association kinetics of EB to the MT plus end, SxIP-EB binding, inherent kinetics between the iLID lit and dark state conformations in the presence and absence of blue light, and the association kinetics of the ssrA sequence with SspB (Fig. 3B (inset)). We defined the apparent off-rate as the rate of loss of tgRFP-SspB fluoresence from MT plus ends post blue light exposure after peak recruitment of tgRFP-SspB. *Drosophila* S2 cells co-transfected with SKIP-LZ-iLID and tgRFP-SspB were exposed to blue light for 200 msec to activate tgRFP-SspB plus end tracking. Time lapse images to monitor tgRFP-SspB localization were initiated at 3 second intervals and blue light exposure was terminated, enabling the iLID system to return to the dark state (Fig. 3A and B; and Video 3). Whole cell tgRFP-SspB comet intensities were quantitated over time. tgRFP-SspB MT plus end tracking activity observed in the frame following blue light activation showed an initial increase in signal/plateau over a 30 sec timeframe and then decayed with an apparent half life of 25.8 ± 1.3 seconds (Fig. 3B). The apparent MT plus end off rate for the SxIP iLID system indicates that it can deactivate within a timeframe appropriate for probing many cytoskeletal activities. This apparent decay rate is on par with the reported *in vitro* iLID decay rate (18 ± 2 sec.) and cellular iLID decay rates in systems designed for membrane localization (52.5 ± 2 sec.) and mitochondrial localization (23.2 ± 1.5 sec.) (Hallett et al., 2016). Once tgRFP-SspB dissociated from MTs, it could again be recruited to MT plus ends, demonstrating the system’s efficacy for multiple, reiterative recruitment cycles (Fig. 3C and Video 4).

**Figure 3.**
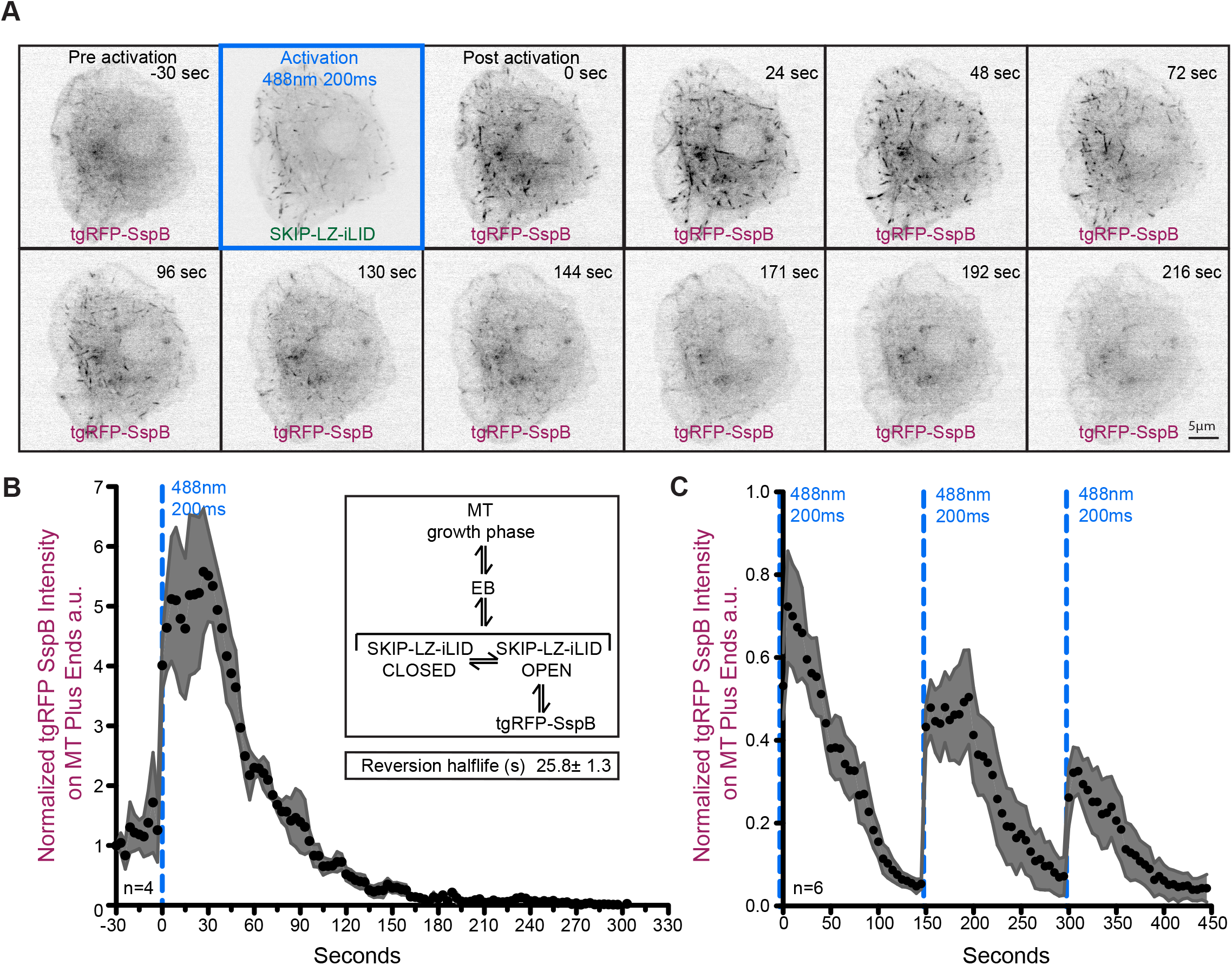
Figure 3. **Dynamics of blue light-dependent tgRFP-SspB MT plus end recruitment and dissociation. (A)** Montage of tgRFP-SspB pre activation and post activation in a *Drosophila* S2 cell co-transfected with eGFP-SKIP-LZ-iLID. Time in seconds is relative to t=0 at which time the cell was activated with a single 200 msec pulse of 488 nm light. The montage shows blue light-dependent tgRFP-SspB recruitment to MT plus ends as well as subsequent dissociation over time (see accompanying Video 3). Scale bar, 5 μm. **(B)** Plot displaying the apparent cellular kinetics of tgRFP-SspB association to, and disassociation from, MT plus ends over time. The system was activated at t=0 with a single pulse of 488 nm light. n=4 cells. In cells co-transfected with SKIP-LZ-iLID, tgRFP-SspB immediately associates with MT plus ends, reaches maximal recruitment within 30 seconds post activation, and dissociates within 90 seconds. Inset shows the general kinetic steps that enable tgRFP-SspB MT plus end association, the convolution of which yields the apparent kinetics observed. The apparent half life of tgRFP-SspB on the MT plus end post activation is 25.8 ± 1.3 sec. **(C)** Multiple rounds of activation of cells co-transfected with SKIP-LZ-iLID and tgRFP-SspB show the ability of tgRFP-SspB to be recruited to MT plus ends multiple times. Cells were activated at t=0, t=150 and t=300 seconds with a single pulse of 488 nm light (see accompanying Video 4). n= 6 cells. For (B) and (C) black points and grey area represent the mean and SEM respectively.

### Cross-linking MT plus ends to the F-actin network decreases MT growth rates and increases the area of the cell void of MTs in *Drosophila* S2 cells

Cytoskeletal networks are cross-linked and their dynamic actions integrated and regulated. How cross-linking dictates downstream cell morphological changes is poorly understood. To understand how coupling of the F-actin and MT cytoskeleton networks affects MTs and MT network architecture, we examined how the optogenetic recruitment of an F-actin binding domain to MT plus ends would affect MT dynamics. Spectaplakins are large MT-F-actin cross-linking proteins with N-terminal tandem calponin homology domains (CH-CH) that bind F-actin and a C-terminal MT-binding module that includes a MT-binding GAR domain and an EB-binding SxIP motif that collectively localize spectraplakins to MT plus ends (Fig. 4A). We fused the tandem CH-CH domain from the sole *Drosophila* spectraplakin Shot to tgRFP-SspB (Fig. 4A) and cotransfected *Drosophila* S2 cells with CH-CH-tgRFP-SspB and SKIP-LZ-iLID. Pre-activation, the CH-CH-tgRFP-SspB construct demonstrated retrograde flow-like behavior, indicative of coupling to the dynamic F-actin lamellar network (Fig 4A’ and Video 5). We next pulsed cells with blue light, and monitored eGFP and tgRFP signals. CH-CH-tgRFP-SspB was robustly recruited to MT plus ends upon blue light activation (Fig. 4B and Video 6). We tracked eGFP comets to determine how CH-CH-tgRFP-SspB recruitment affected MT plus end comet velocities. Surprisingly, MT plus end comet velocities decreased dramatically from 11.6 ± 1.9 μm/min (tgRFP-SspB control) to 7.3 ± 2.1 μm/min when CH-CH-tgRFP-SspB was recruited to MT plus ends (Fig. 4D). To determine the long-term effect of MT-actin cross-linking on MT dynamics we co-transfected cells with the constitutively lit mimetic, SKIP-LZ-iLID^Lit^, and CH-CH-tgRFP-SspB (Fig. 4C and Video 6), induced expression of the iLID construct, and analyzed cells 24-30 hours post induction. MT comet velocities in cells with long-term cross-linking (24-30 hours) mimicked that of the optogenetically cross-linked networks (12.2 ± 2.0 μm/min control versus 6.5 ± 2.2 μm/min crosslinked) (Fig. 4B-D). These data suggest that within a short time frame (3 minutes) MT comet velocities are slowed and remain slowed to the same degree as after a 24-30 hour period of constitutive cross-linking.

**Figure 4.**
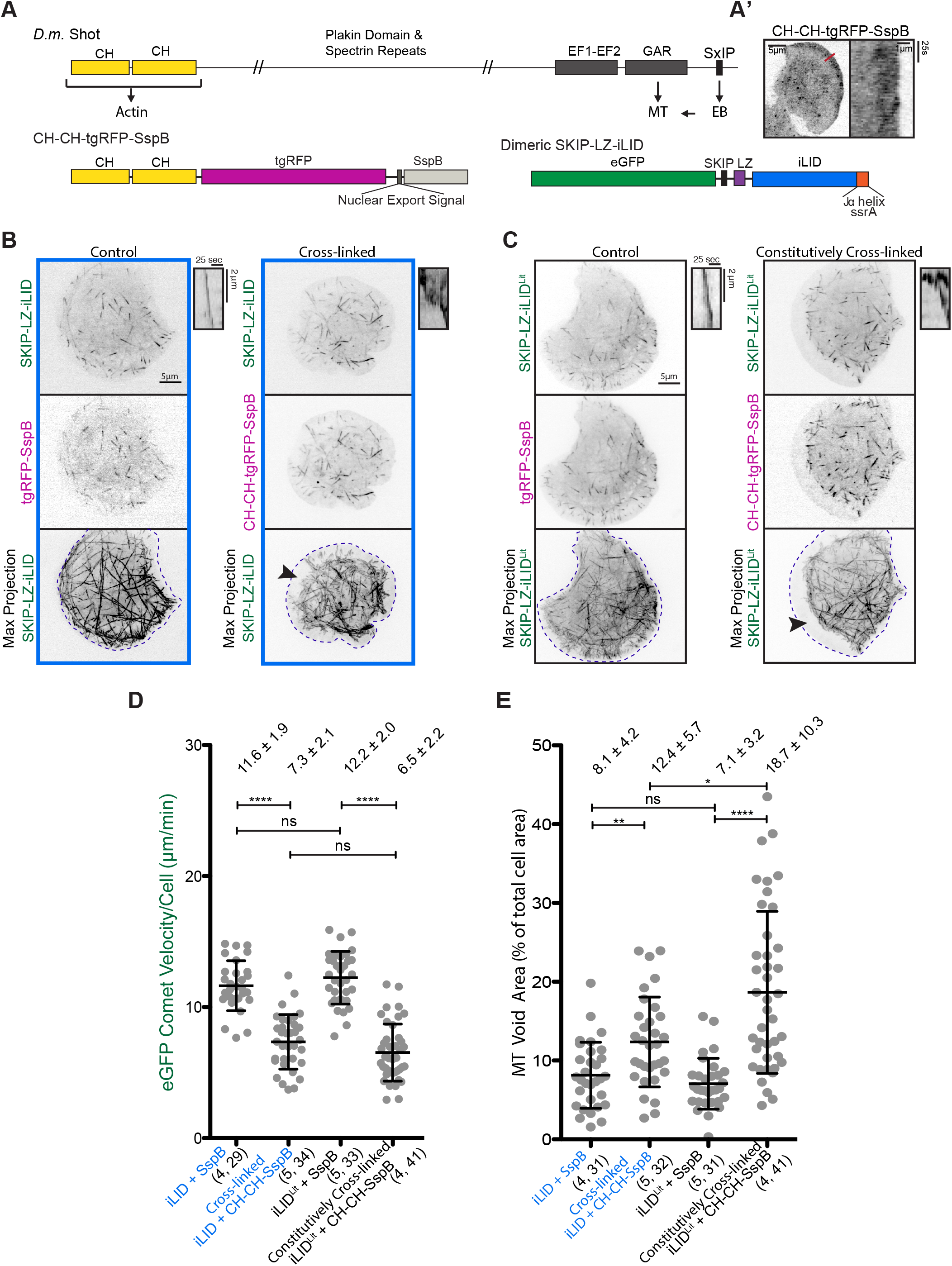
Figure 4. **Optogenetically-induced cytoskeletal cross-linking decreases MT comet velocities and increases MT void lamellar area. (A)** Schematic of the *Drosophila* spectraplakin family member Shot (*D.m.* Shot). Shot contains tandem N-terminal CH domains that bind F-actin, a C-terminal GAR domain that binds MTs, and a SxIP motif that confers MT plus end localization. We functionally parsed the bivalent F-actin - and MT-binding activity of Shot into the two components of our iLID MT plus end tracking system. Shot’s tandem CH domains were fused to tgRFP-SspB. The SKIP-LZ-iLID construct tracks MT plus ends using a SxIP motif from the mammalian spectraplakin MACF2. (A’) Image of CH-CH-tgRFP-SspB in a transfected *Drosophila* S2 cell (left). Scale bar, 10 μm (see accompanying Video 5). The kymograph at right represents a time-lapse of the red line scan at left, and shows retrograde movement of the CH-CH-tgRFP-SspB construct in the lamellar region. Kymograph scale bars: 1 μm, 25 seconds. **(B)** Representative images of S2 cells co-transfected with eGFP-SKIP-LZ-iLID and either a tgRFP-SspB control, or the F-actin-binding CH-CH-tgRFP-SspB construct, and repeatedly pulsed with blue light (250 msec every 3 seconds at 488 nm). Top and middle panels show single images from the GFP and RFP channels. Scale bar, 5 μm. Kymographs at right show representative eGFP-SKIP-LZ-iLID plus end comets. Kymograph bars, 2 μm, 25 sec. Max projections of 60 frames (total 3 min) are shown below, revealing the area of the cell traversed by eGFP-SKIP-LZ-iLID-containing MT plus ends. Areas void of MT plus ends can be seen in the cross-linked cell (arrowhead). Blue boxes indicate blue light recruitment of tgRFP-SspB constructs. **(C)** Representative images of S2 cells co-transfected with eGFP-SKIP-LZ-iLID^Lit^ and either a tgRFP-SspB control, or the F-actin-binding CH-CH-tgRFP-SspB construct. Top and middle panels show single images from the GFP and RFP channels. Scale bar, 5 μm. Kymographs at right show representative eGFP-SKIP-LZ-iLID^Lit^ plus end comets. Kymograph bars, 2 μm, 25 seconds. Max projections of 60 frames (total 3 minutes) are shown below, revealing the area of the cell traversed by eGFP-SKIP-LZ-iLID^Lit^-containing MT plus ends. Areas void of MT plus ends can be seen in the constitutively cross-linked cell (see accompanying Video 6). **(D)** eGFP comet velocity/cell (μm/min) for control cells (iLID +SspB and iLID^Lit^+SspB), light-activated cross-linked cells (iLID + CH-CH-SspB) and constitutively cross-linked cells (iLI D^Lit^ + CH-CH-SspB). Cells with blue light-activated cross-linking and with constitutively cross-linked MT and F-actin networks show decreased eGFP comet velocities compared to control cells. Numbers in parenthesis indicate: (number of experiments, total number of cells quantified). Line represents the mean, error bars indicate SD. P-values were determined by two-way unpaired Student’s *t* test. **** p<0.0001 **(E)** MT void area as a percentage of the total cell area for control and cross-linked cells. Max projection images of 60 frames (spanning 3 minutes) were used to determine the area of the cell void of MTs (representative image used for quantification are shown in B and C and Video 6). Blue light-activated cross-linked and constitutively cross-linked cells show an increase in the area of the cell void of MTs. Numbers in parenthesis indicate: (number of experiments, total number of cells quantified). Line represents the mean, error bars indicate SD. P-values were determined by two-tailed nonparametric Mann–Whitney *U* test. *p<0.05, **p<0.005, **** p<0.0001

Blue-light induced MT plus end-F-actin cross-linking also resulted in a dramatic exclusion of MT plus ends from the lamellar region (Fig. 4B and C (max projections, see arrowheads) and E; and Video 6). When cells expressing CH-CH-tgRFP-SspB and SKIP-LZ-iLID were continuously activated for 3 minutes, the ability of MTs to enter the F-actin-rich lamellar zone of the cell was compromised. Figure 4C shows maximum intensity projections of 60 frames (3 minutes of activated cross-linking) illustrating the dramatic change in the distribution of MTs within the cell upon cross-linking. The area of the cell void of MTs increased from 8.1 ± 4.2% to 12.4 ± 5.7% (Fig. 4E). This MT void region was further enhanced (18.7 ± 10.3% of the cell’s area) when the SKIP-LZ-iLID^Lit^ construct was used with the CH-CH-tgRFP-SspB construct to induce longterm (24-30 hours) cross-linking. These data suggest that the dynamic branched F-actin network in the lamella immediately engaged growing MT plus ends and continuously restricted their entry into the lamella.

### The effects of MT-F-actin cross-linking are F-actin-dependent

To determine if the decrease in MT comet velocity and the increase in MT-void lamellar area in cross-linked cells were F-actin-dependent, we treated SKIP-LZ-iLID^Lit^-transfected cells with latrunculin A to inhibit F-actin assembly. Titrating latrunculin A significantly reduced the ability of the CH-CH-tgRFP-SspB construct to retard MT growth rates (Fig. 5A and B). In cells in which CH-CH-tgRFP-SspB was constitutively recruited to MT plus ends, the mean MT growth velocity was partially restored to WT levels with 2 nM latrunculin A (8.9 ± 2.7 μm/min) while 2 μM latrunculin resulted in a mean MT growth velocity that exceeded that observed in the control tgRFP-SspB transfected cells (13.9 ± 3.9 μm/min vs. 12.2 ± 1.9 μm/min) (Fig. 5B). This result could potentially be due to liberating MTs from endogenous MT-F-actin cross-linking activity. The ability of the CH-CH-tgRFP-SspB construct to exclude MTs from the lamellar zone was also F-actin-dependent as titrating latrunculin A promoted the ability of MTs to fully occupy the area (particularly the perimeter zone) of the cell (Fig. 5A, max projection). When the MT-void area was quantified in cells in which CH-CH-tgRFP-SspB was constitutively recruited to MT plus ends, 2 nM latrunculin A treatment lead to a slight decrease in the MT-void area compared to DSMO control-treated cells (13.4 ± 5.8 % vs. 17.4 ± 7.3 %) while 2 μM latrunculin A treatment dramatically decreased the MT-void area to 4.1 ± 3.0 %, significantly less than that of SspB control transfected cells treated with DMSO (7.3 ± 3.2 %), suggesting again that endogenous cross-linkers promote MT exclusion from the lamellar zone (Fig. 5C). In support, the MT-void region in SspB control transfected cells was further reduced when treated with 2 μM latrunculin A (4.4 ± 2.9 %, not significantly different from the CH-CH-tgRFP-SspB transfected cells).

**Figure 5.**
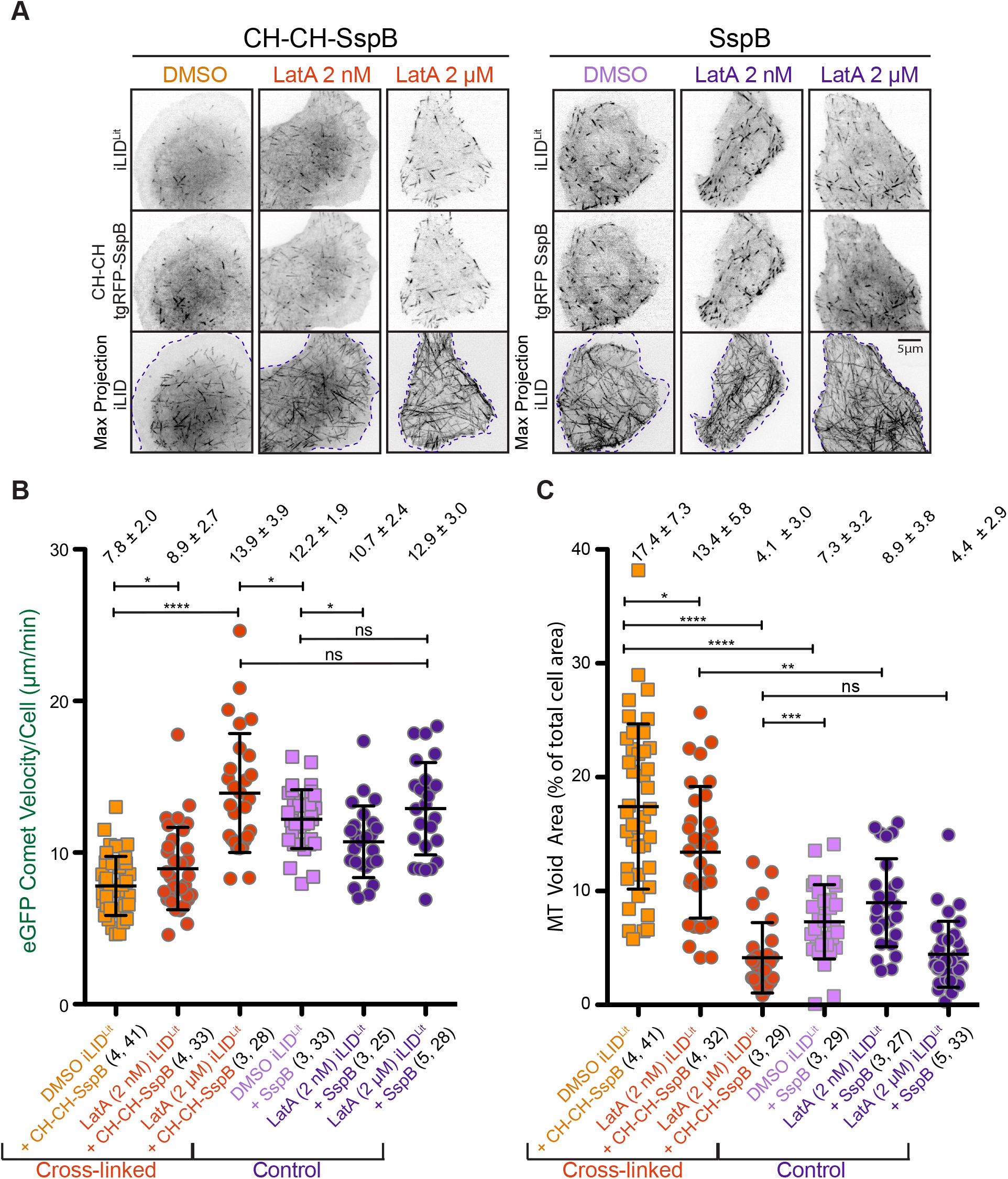
Figure 5. **Optogenetically induced cross-linking decreases MT comet velocities and increases the area void of MT plus ends in an F-actin dependent manner. (A)** Representative images of eGFP-SKIP-LZ-iLID^Lit^ (iLID^Lit^) co-expressed with CH-CH-tgRFP-SspB or tgRFP-SspB in S2 cells treated with DMSO as a control, or Latrunculin A (LatA; 2 nM or 2 μM) for one hour prior to imaging. Top and middle panels show GFP and RFP channels respectively. Bottom panel: max projections of the GFP channel (60 frames collected over 3 min) show that in constitutively cross-linked cells, MT plus ends are excluded from the actin rich lamellar region in DMSO control treated cells, but when the actin network is depolymerized via treatment with LatA, MT plus ends traverse all the way to the cell edge. Scale bar: 5 μm. **(B)** eGFP comet velocity/cell (μm/min) for control cells (iLI D^Lit^ + SspB) and constitutively cross-linked cells (iLI D^Lit^ + CH-CH-SspB) treated with DMSO or LatA (2 nM or 2 μM). Constitutively cross-linked cells treated with LatA show an increase in comet velocities compared to DMSO treated cells. Numbers in parenthesis indicate: (number of experiments, total number of cells quantified). Central lines represent the mean, error bars indicate SD. P-values were determined by two-way unpaired Student’s *t* test. * p<0.05, **** p<0.0001. **(C)** MT void area as a percentage of the total cell area for constitutively cross-linked cells and control cells treated with DMSO, 2 nM LatA, or 2 μM LatA. Max projection images of 60 frames (collected over 3 minutes) were used to determine the area of the cell void of growing MT plus ends (representative image used for quantification shown in A). MT void area decreased in constitutively cross-linked cells with the addition of LatA as compared to control DMSO treatment. Numbers in parenthesis indicate: (number of experiments, total number of cells quantified). Central lines represent the mean, error bars indicate SD. P-values were determined using a two-tailed nonparametric Mann-Whitney *U* test. * p<0.05, ** p<0.005, ***p<0.0005, **** p<0.0001.

### MT-F-actin cross-linking decreases comet velocities and increases void lamellar area in a MT plus end dependent manner

To determine if the decrease in MT comet velocity and increase in the MT-void lamellar area were dependent on MT plus end engagement we co-transfected cells with a CH-CH-tgRFP construct lacking the SspB domain and either SKIP-LZ-iLID or SKIP-LZ-iLID^Lit^ (Fig. 6A). In contrast to the dramatically decreased comet velocities observed when CH-CH-tgRFP-SspB was recruited to MT plus ends (7.3 ± 2.1μm/min in SKIP-LZ-iLID-expressing cells and 6.5 ± 2.2 μm/min in SKIP-LZ-iLID^Lit^ expressing cells), expressing CH-CH-tgRFP yielded MT comet velocities of 9.5 ± 2.2 μm/min and 9.8 ± 1.4 μm/min in SKIP-LZ-iLID and SKIP-LZ-iLID^Lit^ expressing cells, respectively (Fig. 6B). This indicates that recruiting the F-actin-binding CH-CH domain to MT plus ends effectively retards MT comet velocities. Of note, expressing the CH-CH-tgRFP construct that lacked iLID binding capabilities did yield slightly reduce MT comet velocities (9.5 ± 2.2 μm/min tgRFP-CH-CH vs. 11.6 ± 1.9 μm/min tgRFP-SspB control) which may be the result of CH-CH-dependent effects on the stability and/or density of the F-actin network, which in turn could affect MT comet velocity. Although CH-CH-tgRFP expression partially reduced MT comet velocities, MTs were still able to enter the lamellar region (Fig. 6A, max projection) which contrasts with the MT exclusion observed when CH-CH-tgRFP-SspB was recruited to MT plus ends. In cells expressing CH-CH-tgRFP and SKIP-LZ-iLID or SKIP-LZ-iLID^Lit^, the MT-void area was not significantly different than in control cells (Fig. 6C).

**Figure 6.**
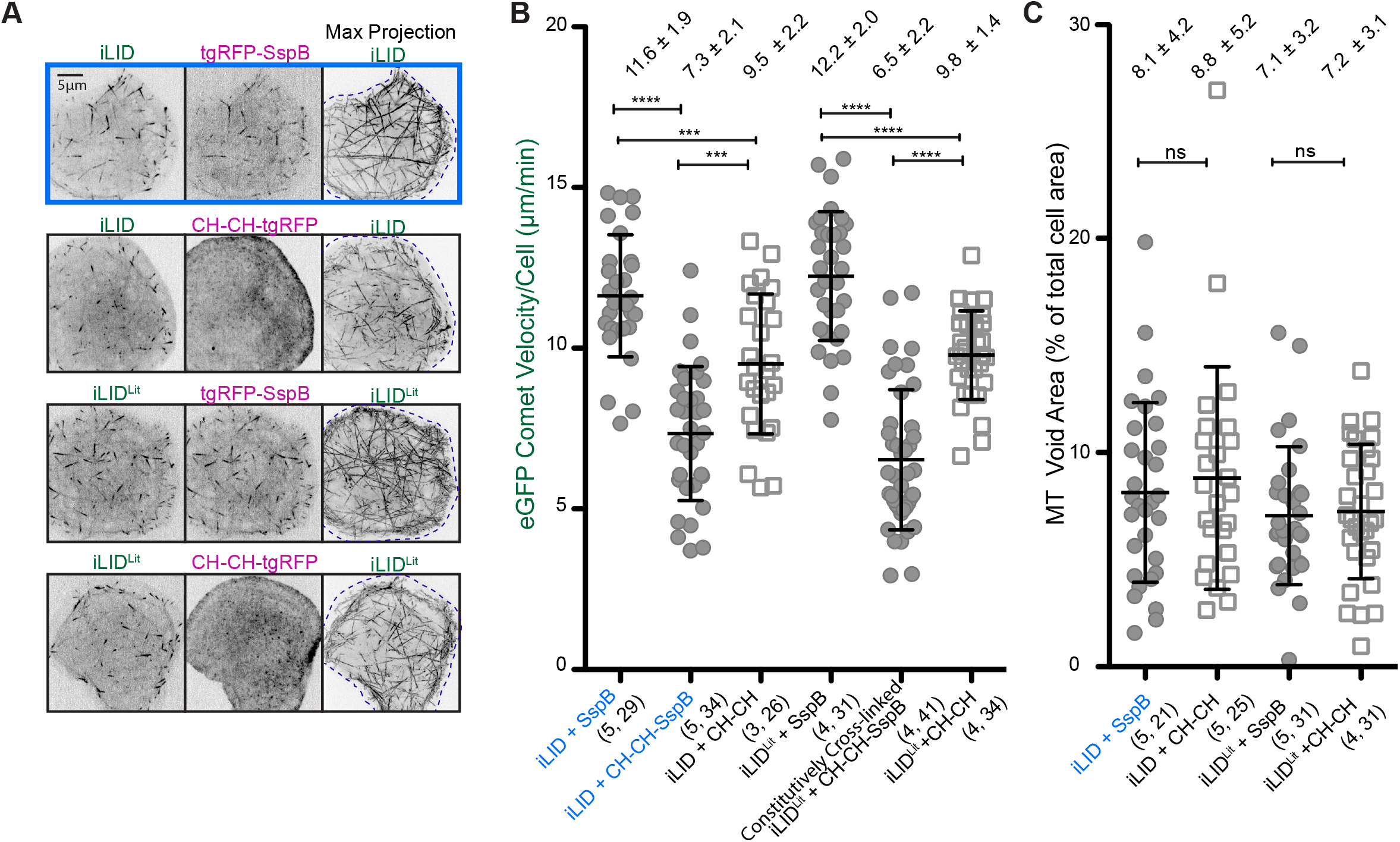
Figure 6. **CH-CH MT plus end recruitment is required to decrease MT comet velocities and generate a peripheral zone where MTs are excluded. (A)** Representative still images of SKIP-LZ-iLID or SKIP-LZ-iLID^Lit^ co-expressed with CH-CH-tgRFP or tgRFP-SspB in cells, showing GFP and RFP channels. Panels at right: max projections of 60 frames (collected over 3 minutes) showing the area traversed by MT plus ends. Scale bar: 5 μm. **(B)** eGFP comet velocity/cell (μm/min) for cross-linked control cells (iLID + SspB activated with blue light and iLI D^Lit^ + SspB) and cells transfected with a CH-CH-tgRFP construct that binds F-actin, but lacks the SspB domain that mediates cross-linking to MT plus ends (iLID + CH-CH and iLI D^Lit^ + CH-CH). Cells in which the CH-CH construct engages the actin network but cannot engage MT plus ends yields MT comet velocities that are near to control, non-cross-linked velocities. Numbers in parenthesis indicate: (number of experiments, total number of cells quantified). Central lines represent the mean, error bars indicate SD. P-values were determined by two-way unpaired Student’s *t* test. ***p<0.0005, **** p<0.0001. **(C)** Area void of MT plus ends as a percentage of the total cell area for control cells (iLID + SspB activated with blue light and iLID^Lit^ + SspB) and cells in which the CH-CH construct engages the actin network, but is not cross-linked to MT plus ends (iLID + CH-CH and iLI D^Lit^ + CH-CH). Max projection images of 60 frames (collected over 3 minutes) were used to determine the area of the cell void of MT plus ends (representative image used for quantification shown in A). The MT void area does not change significantly if MT plus ends do not engage the CH-CH construct. Numbers in parenthesis indicate (number of experiments, total number of cells quantified). Central lines represent the mean, error bars indicate SD. P-values were determined using a two-tailed nonparametric Mann-Whitney *U* test.

These data show that cross-linking of the F-actin and MT networks in *Drosophila* S2 cells leads to decreased MT comet velocities and an increase in the area of the cell void of MTs in an F-actin and MT plus end-dependent manner. Cross-linking rapidly stalls MTs and prevents entry into the peripheral region of the cell.

## Discussion

Advances in cellular optogenetics have allowed for spatial and temporal control of many biological processes, enabling researchers to test the role of specific proteins or protein domains during different phases of the cell cycle and in different cellular locations. Here, we generated a novel optogenetic tool, SxIP-iLID, which can recruit proteins of interest to MT plus ends. This tool provides broad utility to probe the spatiotemporal role of diverse MT effectors at cellular and organismal levels. A library of SxIP-iLID constructs has been generated that offers a range of MT plus end tracking activities that can be selectively utilized to match the physiological requirements of a specific cytoskeletal processes. In addition, the dimeric SKIP-LZ-iLID construct would enable avidity-based recruitment of multimeric SspB-tagged constructs. The SxIP-iLID system also serves as a generalizable platform that can incorporate recently characterized iLID mutations that affect activation and factor recruitment kinetics (Zimmerman et al., 2016). The system is superior to chemical-induced heterodimerization as reversion is rapid and does not require drug washout, the system can be repeatedly re-activated, and microscopes that can activate iLID with 405-488 nm light and can monitor a fluorescent reporter (e.g. RFP/mCherry) in another channel are readily available to the community.

As a proof of principal we used the SxIP-iLID system to examine the effects of crosslinking the F-actin and MT networks using a minimalist spectraplakin analog. While it is known that spectraplakins play critical roles in cytoskeletal cross-linking, how cross-linking activity affects the morphology and dynamics of the MT network is poorly understood. We fused the N-terminal CH, actin binding domains from the *Drosophila* protein Shot to SspB, generating a light inducible cross-linking system when used with the SKIP-LZ-iLID construct. Light-induced MT-F-actin crosslinking rapidly decreased MT comet velocities and increased the cellular area void of MTs. Whether the decreased MT comet velocity is due to diminished MT polymerization rates or MT sliding remains to be determined. Our results suggest that cross-linking in cells may mechanically stall/slow MT growth and/or entry into the F-actin rich lamella. This may reflect the role of MT-F-actin cross-linkers in focal adhesion turnover during cell migration.

## Materials and methods

### Molecular biology

The iLID micro domain (Guntas et al., 2015) was amplified and sub-cloned using the Gateway TopoD pEntr system (Invitrogen) into an ampicillin-selectable backbone containing a methallothionein promoter and a N-terminal GFP (pMTeGFP). The minimal MT plus end tracking motif from MACF2 (Slep et al., 2005; Honnappa et al., 2009) followed by a two-stranded leucine zipper (LZ) coiled-coil sequence corresponding to GCN4-p1 (Steinmetz et al., 2007) was PCR amplified with primers containing restriction enzyme sites, and cloned into the pMTeGFP-iLID backbone. The remaining SxIP-iLID constructs were generated via site directed mutagenesis to the pMTeGFP-LZ-SKIP-iLID parent vector. The constitutively lit mutant (I539E of the iLID domain (Harper et al., 2004): SKIP-LZ-iLID^Lit^) was generated using KOD Xtreme site directed mutagenesis following the manufacture’s protocol (Novagen).

tgRFPt-NES-SspB nano (referred to as tgRFP-SspB in this paper) was PCR amplified from a pLL7.0 vector and sub-cloned using the Gateway TopoD pEntr system (Invitrogen) into a final ampicillin-selectable actin promoter backbone (pAW). DNA encoding the tandem CH domains from *Drosophila* Shot was PCR amplified and inserted into the pAW-tgRFPt-NES-SspB nano construct using restriction enzymes. The EB1-GFP expression construct, under control of the EB1 promoter, was constructed as described in Currie et al., 2011.

### Cell culture and transfection

*Drosophila* S2 cells were cultured using the standard protocol described in Rogers and Rogers, 2008. In short, S2 cells (*Drosophila* Genomics Resource Center, Bloomington, IN) were grown in Sf900II SFM (Gibco) supplemented with 1x antibiotic-antimycotic (Invitrogen). To express the desired constructs, 2.5 x 10^5^ cells were plated in 12 well dishes and transfected with 1.5 mg of DNA using FuGENE HD (Promega) following the manufacturer’s protocol. Expression of pMT constructs was induced using 50 μM of CuSO_4_ 24 hours post transfection and 20-30 hours prior to imaging.

### Live-cell imaging

S2 cells were seeded onto 0.5 mg/ml conconavlin A (ConA)-treated glass bottom imaging dishes in 1 ml of Schneider’s *Drosophila* media (Gibco) supplemented with 1x antibiotic-antimycotic (Invitrogen) and 10% fetal bovine serum (Gibco) and allowed to spread for 2 hours prior to imaging. Imaging dishes were prepared by attaching glass coverslips to drilled 35 mm tissue culture dishes using UV curable glue (Norland Products). Time-lapse imaging was performed on a VT-HAWK confocal system (Visitech) using an inverted microscope (Nikon Ti) equipped with a 100x/1.45 N.A. objective lens driven by VoxCell software (VisiTech). Images were captured with a Flash 4.0 camera (Hamamatsu). To assess comet velocities in cells exclusively transfected with either EB1-GFP or an eGFP-SxIP-iLID construct, time lapse images were acquired every 2 seconds using 488 nm excitation. To assess comet velocities in cells co-transfected with an eGFP-SxIP-iLID construct and a tgRFP-SspB construct, cells were imaged every 3 seconds using alternating 561 nm and 488 nm excitation. To determine the apparent off rate of the system, images were acquired every 3 seconds using 561 nm excitation for 10 frames, a single pulse of 488 nm light was then applied to induce recruitment of the tgRFP-SspB construct to MT plus ends, then excitation at 561 nm resumed and frames were captured every 3 seconds over the course of 3 minutes to record the re-localization of the SspB-tgRFP construct to the cytoplasm. To analyze tgRFP-SspB behavior over multiple rounds of activation, single pulses of 488 nm light were applied to induce recruitment of the tgRFP-SspB construct to MT plus ends at t=0, t=150 and t=300 sec. After each 488 nm pulse, frames were captured every 5 seconds for 2.5 minutes using 561 nm excitation. Images were processed using Fiji (Schindelin et al., 2012).

### Comet velocity, intensity analysis, and kinetic rate analysis

Comet velocities were determined using the MTrackJ plugin in Fiji (Meijering et al., 2012, Schindelin et al., 2012). Twelve MT plus ends were hand tracked per cell for 10 or more consecutive frames. These tracks were then averaged to give the average comet velocity per cell. Fiji was used to determine the total cell fluorescence intensity/area. The cell of interest was selected using the freeform selection tool and an area next to the cell was selected for background intensity measurement. The integrated density and area of these selections was measured. The total cell intensity was then calculated as: [Integrated density – (Area of selected cell * Mean fluorescence of background readings per area)]. This was then reported as total cell intensity/area of the selected cell. To determine the comet/cytoplasmic intensity ratio, 10 comets were analyzed per cell. Using the freeform selection tool the region of the comet was outlined and the integrated density and the area of each comet was determined. As the cytoplasmic intensity varies throughout the cell, the region surrounding each comet was selected to determine the respective mean cytoplasmic intensity per area by selecting a rectangle encompassing the entire comet. The integrated density and the area of this rectangle were determined. To determine the integrated density of the cytoplasmic region, the integrated density of the comet was subtracted from the integrated density of the rectangle. To determine the area of the cytoplasmic region, the area of the comet was subtracted from the area of the rectangle. The integrated density per area for both the comet and the cytoplasmic region were then determined and corrected for non-cellular background noise. The ratio of the integrated density per area of the comet versus the cytoplasm was then calculated and reported as comet/cytoplasm intensity. Therefore, a value of 1 would represent a cell in which the comet and the cytoplasm have the same mean intensity per area. To determine the tgRFP-SspB intensity on MT plus ends for the plots of apparent cellular kinetics for single and multiple rounds of activation, a threshold was applied to encompass the tgRFP-SspB signal at MT plus ends using the first frame post activation (t=0 seconds). This threshold was then applied to all frames, pre and post activation and the intensity was calculated as [Integrated Density – (Area of threshold region* Mean fluorescence per area of the non cellular background)]. For the plot of apparent cellular kinetics, each individual cell plot was normalized to the first time point (t=-30 seconds). The apparent reversion half life was determined by fitting points 22-111 (t=33-300 seconds) which correspond to the peak activation through the last data point. The curve fit was generated using Prism (GraphPad) using one-phase decay with least squares fit. For the plot of multiple rounds of activation each individual cell plot was normalized to the value of peak intensity after the first pulse of blue light.

### Statistical analyses

eGFP comet velocity data were analyzed using a two-way unpaired Student’s *t* test (GraphPad Prism). MT void area data were analyzed using a two-tailed nonparametric Mann-Whitney *U* test (GraphPad Prism). * p<0.05, ** p<0.005, ***p<0.0005, **** p<0.0001.

### Online supplemental material

Fig. S1 shows that the amount of a given eGFP-SxIP-iLID construct either in the cell or on the MT plus end does not strongly correlate with comet velocity.

## Acknowledgments

We thank Seth Zimmerman for the pLL7.0 tgRFPt-NES-SspB vector. We thank Derek Applewhite and Thomas Lane for insightful discussions, and Mark Peifer and Alakananda Das for comments regarding this manuscript. This work was supported by the National Institutes of Health grant RO1GM094415 and March of Dimes grant FY11-434 to K.C. Slep, R.C. Adikes was supported by National Institutes of Health grant F31-GM116476 for this work.

## Author Contributions

Conceptualization: R.C. Adikes, R.A. Hallett, B. Kulhman, K.C. Slep

Data curation: R.C. Adikes

Formal analysis: R.C. Adikes, B.F. Saway, K.C. Slep

Funding acquisition: R.C. Adikes, K.C. Slep

Investigation: R.C. Adikes, B.F. Saway

Methodology: R.C. Adikes, K.C. Slep

Project administration: K.C. Slep

Resources: B.Kulhman, K.C. Slep

Supervision: K.C. Slep

Validation: R.C. Adikes

Visualization: R.C. Adikes

Writing – original draft: R.C. Adikes

Writing – review & editing: R.C. Adikes, B. Kulhman, K.C. Slep The authors declare no competing financial interests.

## Footnotes

Abbreviations used:

MT: Microtubules
LIDs: Light Inducible Dimers
LOV: Light-Oxygen-Voltage
iLID: improved Light Inducible Dimer
EB: End Binding protein
LZ: Leucine Zipper
CH: Calponin Homology
LatA: Latrunculin A

---

**Supplemental Figure1.**
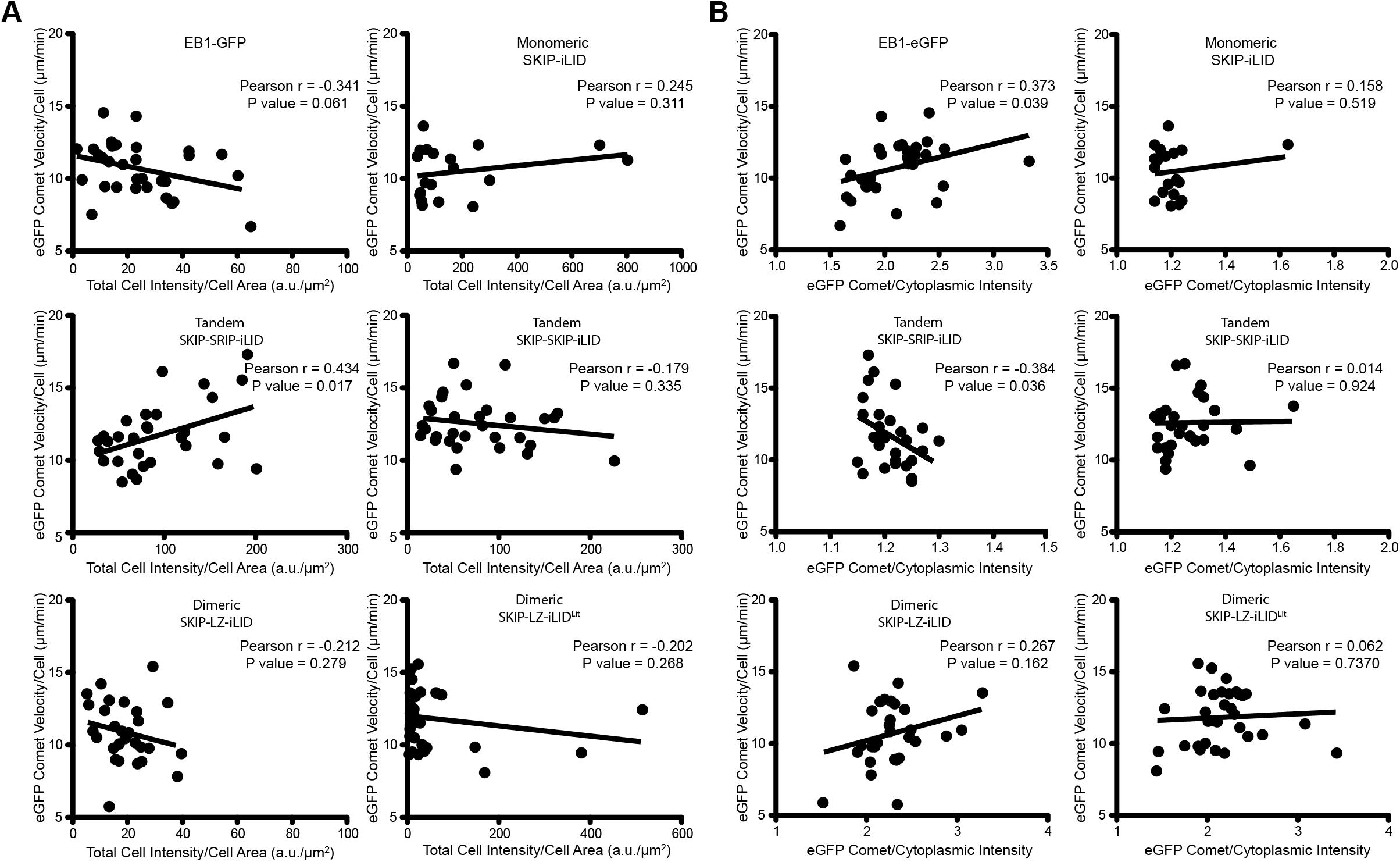
Supplemental Figure 1. **The amount of eGFP-SxIP-iLID in the cell or on the MT plus end does not strongly correlate with comet velocity or comet length. (A)** The total cell intensity/area (a.u./μm^2^ ) was determined for cells transfected with eGFP-SxIP-iLID constructs or EB1-GFP and plotted against the average eGFP comet velocity for each cell. To assess the strength of the correlation, a Pearson’s correlation coefficient was determined. Using this statistical analysis, no correlations we detected except for the SKIP-SRIP-iLID construct which yielded a p-value of less than 0.05, but greater than 0.005. **(B)** The eGFP comet/cytoplasmic intensity plotted against the average eGFP comet velocity for each cell. To assess the strength of the correlation, a Pearson’s correlation coefficient was determined. Using this statistical analysis, no correlations were detected except for the SKIP-SRIP-iLID construct which yielded a p-value of less than 0.05, but greater than 0.01.

## Video legends

Video 1. **eGFP SxIP-iLID constructs in *Drosophila* S2 cells**. *Drosophila* S2 cells were transfected with eGFP-SxIP-iLID constructs 48 hours prior to imaging; expression of the construct was induced 24-30 hours prior to imaging. Images were acquired using a confocal microscope every 2 seconds using 488 nm excitation. Video rate: 5 fps. Corresponding figure: Figure 1.

Video 2. **Photoactivated SxIP-iLID constructs rapidly recruit tgRFP-SspB to MT plus ends.** *Drosophila* S2 cells were transfected with eGFP-SxIP-iLID and tgRFP-SspB constructs 48 hours prior to imaging; expression of the eGFP-SxIP-iLID construct was induced 24-30 hours prior to imaging. Pre-activation tgRFP-SspB is diffuse in the cytoplasm. Upon activation with 488 nm light tgRFP-SspB is rapidly recruited to MT plus ends. Continuous activation with 488 nm light was used to keep tgRFP-SspB at MT plus ends. eGFP-SKIP-LZ-iLID shows the most robust recruitment of tgRFP-SspB upon blue light activation. Cells were imaged on a confocal microscope, images were acquired every 3 seconds using alternating 561 nm and 488 nm excitation. Video rate: 5 fps. Corresponding figure: Figure 2.

Video 3. **tgRFP-SspB is recruited to MT plus ends by SKIP-LZ-iLID after a single pulse of blue light and then rapidly dissociates.** *Drosophila* S2 cells were transfected with eGFP-SKIP-LZ-iLID and tgRFP-SspB constructs 48 hours prior to imaging; expression of the eGFP-SKIP-LZ-iLID construct was induced 24-30 hours prior to imaging. Pre-activation, tgRFP-SspB is diffuse in the cytoplasm. Upon activation with 488 nm light tgRFP-SspB is rapidly recruited to MT plus ends, then quickly dissociates from MT plus ends and returns to the cytoplasm. The cell was imaged on a confocal microscope. Images were acquired every 3 seconds using 561 nm excitation. The cell was activated with a single 200 msec pulse of 488 nm light at t=0, marked by the blue rectangle. Video rate: 5 fps. Corresponding figure: Figure 3.

Video 4. **Multiple rounds of activation of cells co-transfected with eGFP-SKIP-LZ-iLID and tgRFP-SspB show the ability of tgRFP-SspB to be recruited to MT plus ends multiple times.** The *Drosophila* S2 cell was transfected with eGFP-SKIP-LZ-iLID and tgRFP-SspB 48 hours prior to imaging. Expression of the eGFP-SKIP-LZ-iLID construct was induced 24-30 hours prior to imaging. The cell was imaged on a confocal microscope. Images were acquired every 5 seconds using 561 nm excitation. The cell was activated with a single 200 msec pulse of 488 nm light at t=0, t=150, and t=300 seconds, each corresponding frame marked with a blue rectangle. Video rate: 5 fps. Corresponding figure: Figure 3.

Video 6. **Optogenetically-induced cytoskeletal cross-linking decreases MT comet velocities and increases MT void lamellar area.** *Drosophila* S2 cells were transfected with eGFP-SKIP-LZ-iLID and tgRFP-SspB constructs 48 hours prior to imaging. Expression of the eGFP-SKIP-LZ-iLID construct was induced 24-30 hours prior to imaging. The cells were imaged on a confocal microscope. Images were acquired every 3 seconds using alternating 561 nm and 488 nm excitation. Video rate: 5 fps. Corresponding figure: Figure 4.

